# Global changes in patterning, splicing and primate specific lncRNAs in autism brain

**DOI:** 10.1101/077057

**Authors:** Neelroop N. Parikshak, Vivek Swarup, T. Grant Belgard, Michael Gandal, Manuel Irimia, Virpi Leppa, Jennifer K. Lowe, Robert Johnson, Benjamin J. Blencowe, Steve Horvath, Daniel H. Geschwind

## Abstract

We apply transcriptome-wide RNA sequencing in postmortem autism spectrum disorder (ASD) brain and controls and identify convergent alterations in the noncoding transcriptome, including primate specific lncRNA, and transcript splicing in ASD cerebral cortex, but not cerebellum. We characterize an attenuation of patterning between frontal and temporal cortex in ASD and identify *SOX5*, a transcription factor involved in cortical neuron fate specification, as a likely driver of this pattern. We further show that a genetically defined subtype of ASD, Duplication 15q Syndrome, shares the core transcriptomic signature of idiopathic ASD, indicating that observed molecular convergence in autism brain is the likely consequence of manifold genetic alterations. Using co-expression network analysis, we show that diverse forms of genetic risk for ASD affect convergent, independently replicated, biological pathways and provide an unprecedented resource for understanding the molecular alterations associated with ASD in humans.

---

Autism spectrum disorder (ASD) is a neurodevelopmental syndrome characterized by deficits in social communication and mental flexibility^1^. Genetic risk factors contribute substantially to ASD risk, and recent studies support the potential contribution of more than a thousand genes to ASD risk^2-4^. However, given the shared cognitive and behavioral features across the autism spectrum, one hypothesis is that diverse risk factors may converge on common molecular, cellular, and circuit level pathways to result in the shared phenotype^5,6^. Analysis of the transcriptome has been used to identify common molecular pathways in the cerebral cortex (CTX) from postmortem human brain tissue in individuals with ASD^7-11^. However, all transcriptomic studies in ASD to date have been limited to evaluating highly expressed mRNAs corresponding to protein coding genes. Moreover, most lack rigorous replication and do not assess gene expression patterns across brain regions.

We used rRNA-depleted RNA-seq (Methods) to evaluate transcriptomes from a large set of ASD and control (CTL) brain samples including neocortex (frontal and temporal) and cerebellum across 79 individuals (46 ASD, 33 CTL, 205 samples, Extended Data Fig. 1a-e, Supplementary Table 1). We first compared differential gene expression (DGE) between ASD and CTL individuals in CTX from a previously published^7^ microarray study against new, independent gene expression profiles from RNA-seq to evaluate global reproducibility of DGE in ASD. We found a high degree of replication of DGE fold changes between the sample sets, despite evaluation on different gene expression platforms (fold changes at P < 0.05 in previously evaluated data correlate with new data with R^2^ = 0.60, Extended Data Fig. 1f). We observed a much weaker overall signal and replication in cerebellum (R^2^ = 0.033, Extended Data Fig. 1g). These analyses confirm the existence of a reproducible DGE signature in ASD CTX across different platforms and in independent samples.

We next combined samples from all individuals with idiopathic ASD into a covariate-matched “ASD Discovery Set” (Extended Data Fig. 1h) for CTX (106 samples, 26 ASD, 33 CTL individuals) and held out remaining samples for replication (“ASD Replication Set”, Methods). For DGE analysis, we used a linear mixed effects model that accounts for biological and technical covariates (Methods) to identify 1156 genes differentially expressed in ASD CTX, 582 increased and 574 decreased (Benjamini-Hochberg FDR ≤ 0.05, Supplementary Table 2). Importantly, DGE analysis with additional covariates or different assumptions about the distribution of the data and test statistics yielded similar results (Extended Data Fig. 2a). Additionally this DGE signature clusters over two-thirds of ASD samples together and this clustering is not related to confounding factors such as cortical region, age, sex, and RNA quality (Figure 1a, Extended Data Fig. 2b). The most significantly down-regulated gene was *PVALB* (fold change = 0.53, FDR ≤ 0.05), a marker for GABAergic interneurons. *SST*, a marker for a different subpopulation of GABAergic interneurons, is also among the most downregulated (fold change = 0.61, FDR ≤ 0.05). Other down-regulated genes at FDR ≤ 0.05 include *NEUROD6*, involved in neuronal differentiation (fold change = 0.60), multiple ion channels, and *KDM5D*, a lysine demethylase (fold change = 0.66). In contrast, members of the complement cascade implicated in microglial-neuronal interactions (*C4A*, fold change = 1.94; *C1QB*, fold change = 1.65; both FDR ≤ 0.05) are upregulated in ASD CTX. Gene Ontology (GO) term enrichment analysis further supports the involvement of pathways implicated by these genes (Figure 1b), confirming previous findings^7^. Moreover, the upregulated set is enriched for astrocyte and microglia enriched genes, and the down-regulated set is enriched for synaptic genes (Extended Data Fig. 2c), consistent with previous observations^7,11^.

**Figure 1.**
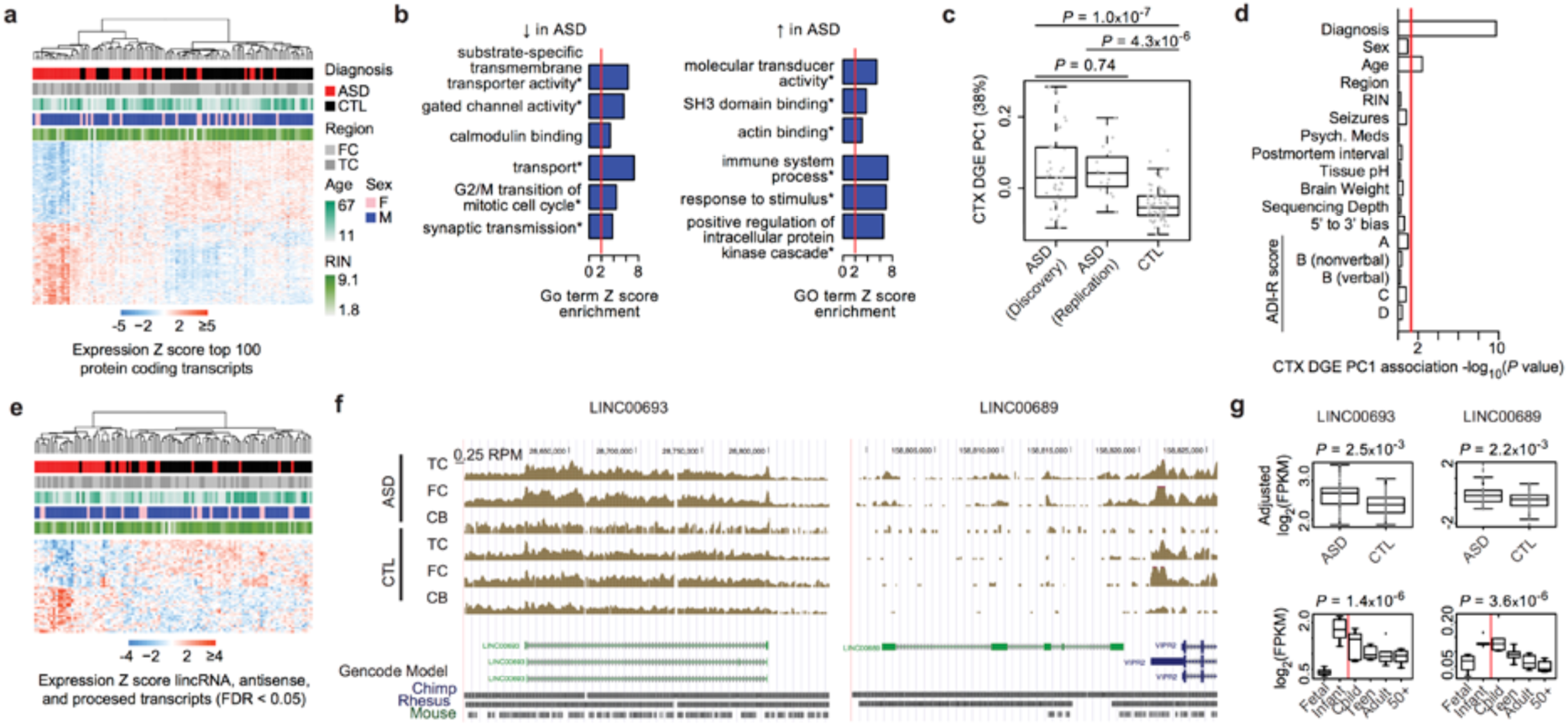
Transcriptome-wide differential gene expression in ASD. a, Average linkage hierarchical clustering of samples in the ASD Discovery Set using the top 100 upregulated and top 100 downregulated protein coding genes. b, Gene Ontology (GO) term enrichment analysis of upregulated and downregulated genes in ASD. *FDR ≤ 0.05 across all GO terms and gene sets. c, 1^st^ principal component of the CTX DGE set (CTX DGE PC1) is able to distinguish ASD and CTL samples, including independent samples from the ASD Replication Set. d, CTX DGE PC1 is primarily associated with diagnosis, and not other factors. e, Average linkage hierarchical clustering of ASD Discovery Set using all lncRNAs in the DGE set. f, UCSC genome browser track displaying reads per million (RPM) in a representative ASD and CTL sample, superimposed over the gene models and sequence conservation for genomic regions including *LINC00693* and *LINC00689*. g, *LINC00693* and *LINC00689* are upregulated across ASD samples and downregulated during frontal cortex development. Abbreviations: FC, frontal cortex; TC, temporal cortex; RIN, RNA integrity number; ADI-R score, Autism Diagnostic Interview Revised score; FPKM, fragments per kilobase million mapped reads.

We next sought to evaluate whether the transcriptional signature identified in the ASD Discovery Set generalizes to the ASD Replication set by assessing the 1^st^ principal component of the DGE set, which summarizes the DGE expression pattern across all cortical samples. The ASD Discovery Set and ASD Replication Set share this pattern, which is significantly different for both sets compared to CTL (Figure 1c). Moreover, this pattern is highly associated with ASD diagnosis, but not other biological factors, technical factors, or scores on sub-domains of an ASD diagnostic tool (Figure 1d). These analyses demonstrate that convergent differences in ASD CTX are reproducible in independent samples and are not related to confounding factors.

We also detected 2715 lncRNAs expressed in cerebral cortex (after careful filtering for high-confidence transcripts, Supplementary Information), of which 62 were significantly dysregulated between ASD and CTL (33 long intergenic RNAs, lincRNAs; 19 antisense transcripts; and 10 processed transcripts at FDR ≤ 0.05). Similar to the protein coding genes, these transcripts’ expression patterns cluster ASD and CTL samples (Figure 1e). Most of these lncRNAs are developmentally regulated^12^, have chromatin states indicative of transcription start sites (TSSs) near their 5´ end in brain^13^, and are identified in other datasets^12,14^ consistent with being valid, functional lncRNAs. Moreover, most (81%) exhibit primate-specific expression patterns in brain^15^ (Supplementary Information). For example, Figure 1f depicts two lincRNAs, *LINC00693* and *LINC00689*, which are typically downregulated during development, yet are upregulated in ASD CTX relative to controls (Figure 1g), which we validated by RT-PCR (Extended Data Fig. 2d). *LINC00693* sequence is present, but poorly conserved in mouse, while *LINC00689* is primate-specific (present in macaque and other primates but not in any other species, Supplementary Information, Extended Data Fig. 3 for additional examples). These data indicate that dysregulation of lncRNAs, many of which are primate-specific and involved in brain development, is an important component of transcriptome dysregulation observed in ASD.

Previous work suggested that alterations in transcript splicing may contribute to transcriptomic changes in ASD^7,16,17^ by evaluating splicing in a targeted manner and pooling samples across individuals^7,16,17^. Given the increased sequencing depth and reduced sequencing bias across transcript length in our dataset, we were able to perform an unbiased genome-wide analysis of differential alternative splicing (AS). We evaluated the percent spliced in (PSI, Extended Data Fig. 4a) for 34,025 AS events in CTX across the ASD Discovery Set, encompassing skipped exons (SE), alternative 5´ splice sites (A5SS), alternative 3´ splice sites (A3SS), and mutually exclusive exons (MXE) using the MATS pipeline^18^ (Supplementary Information). We first asked whether there was a global signal, finding significant enrichment over background (Extended Data Fig. 4b). We identified 1127 events in 833 genes at FDR ≤ 0.5 in CTX (similar to the number of events at uncorrected P < 0.005). Importantly, we obtained similar results with a different splice junction mapping and quantification approach (Extended Data Fig. 4c).

We performed PCR validations with nine AS events from the differential splicing set (*ASTN2, MEF2D, ERC2, MED31, SMARCC2, SYNE1, NRCAM, GRIN1, NCAM*) and found that validated changes in splicing patterns were concordant with RNA-seq (Extended Data Fig. 4d-e), demonstrating that our approach identifies alterations in AS with high specificity. Similar to our observations with lncRNA and DGE, AS changes clustered the samples by diagnosis (Figure 2a). The most significantly different event was the inclusion of an exon in *ASTN2* (ΔPSI = 5.8 indicating a mean of 5.8% difference in inclusion in ASD vs CTL; *P* = 7.8x10^−6^), a gene implicated by copy number variation (CNV) in ASD and other developmental disorders^19^. GO term analysis of the genes implicated by these pathways indicates involvement of biological processes related to neuronal projection, biological adhesion, and morphogenesis (Figure 2b), pathways where alternative isoforms are critical to specifying interactions between protein products. Moreover, the 1^st^ principal component of the cortex differential splicing signature replicates in the ASD Replication Set and is not associated with other biological or technical factors (Figures 2c-d, Extended Data Fig. 5a). Importantly, many splicing alterations occur in genes that are not differentially expressed between ASD and CTL; removing AS events on genes exhibiting even nominal DGE (P < 0.05), still identified a strong difference between ASD and CTL CTX (Extended Data Fig. 5b).

**Figure 2.**
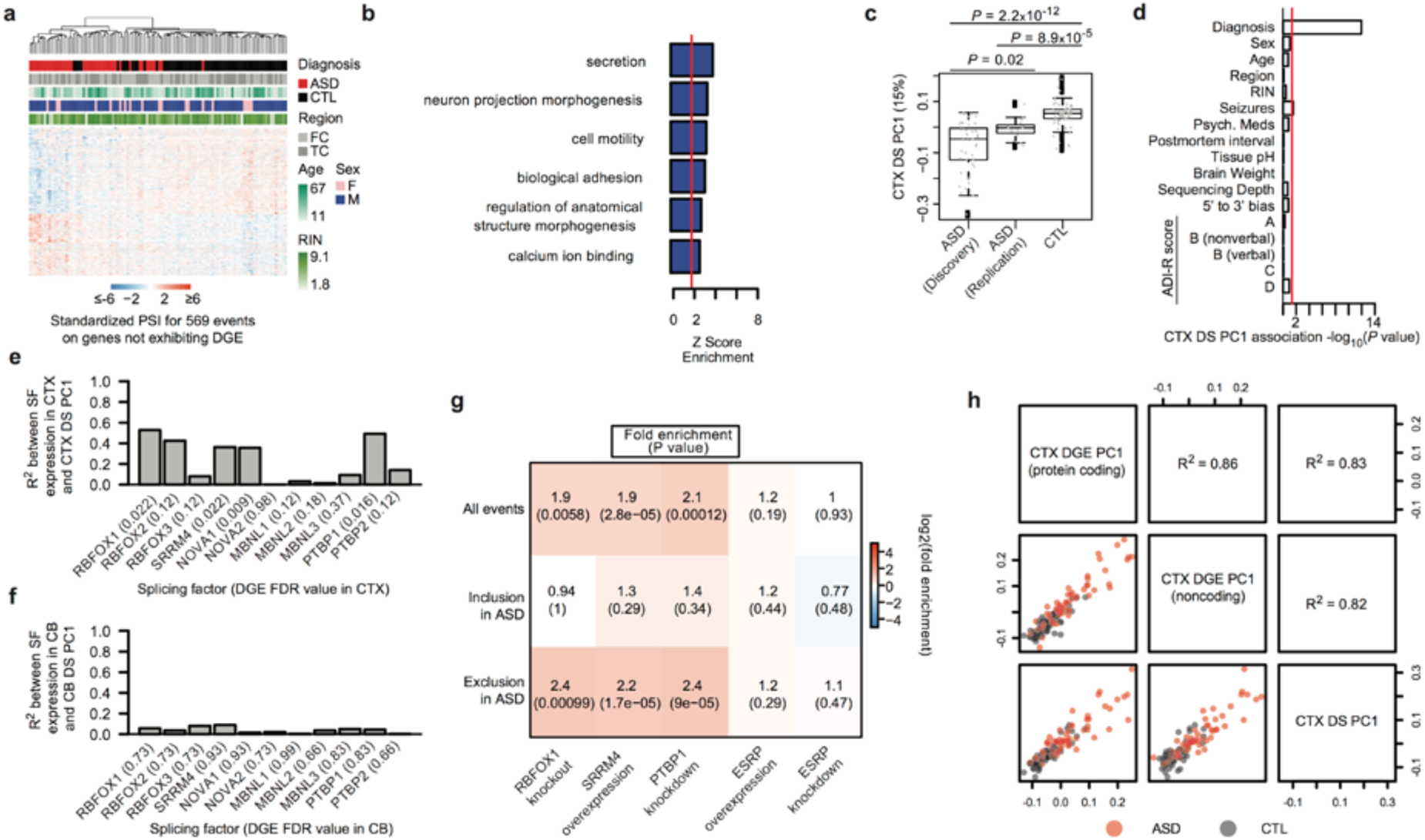
Alteration of alternative splicing in ASD. a, Average linkage hierarchical clustering of ASD discovery set using top 100 differentially included and top 100 differentially excluded exons from the differential splicing (DS) set across the ASD Discovery Set. b, Gene Ontology term enrichment analysis of genes with DS in ASD. c, 1^st^ principal component 1 of the CTX differential alternative splicing set (CTX DS PC1) is able to distinguish ASD and CTL samples using independent samples from the ASD Replication Set. d, CTX DS PC1 is primarily associated with diagnosis, and not other factors. e, Correlation between CTX DS PC1 and gene expression of neuronal splicing factors in CTX. f, Correlation between 1^st^ principal component of cerebellum differential splicing (CB DS PC1) and gene expression of neuronal splicing factors in cerebellum. g, Overlap between DS set and splicing events regulated by splicing factors where experimental data was available. h, Scatterplots and correlations between the 1^st^ principal component across the ASD versus CTL DGE sets for different transcriptome subcategories. Abbreviations: FC, frontal cortex; TC, temporal cortex; RIN, RNA integrity number; ADI-R score, Autism Diagnostic Interview Revised score; FPKM, fragments per kilobase million mapped reads.

A parallel analysis in cerebellum evaluating 32,954 AS events found no differentially regulated events significant at any multiple comparison correction thresholds (Extended Data Fig. 5c, Supplementary Table 3). There was no detectable global overlap between cerebellum and CTX above chance for events significant at P < 0.05 in both comparisons (fold enrichment = 1.1, P = 0.21). This suggests that AS alterations in ASD are largely confined to CTX cell types, consistent with the stronger overall DGE patterns observed in CTX versus cerebellum.

To further explore the underlying biology of AS dysregulation, we tested whether the shared splicing signature in ASD might be a product of perturbations in AS factors known to be important to neural development or preferentially expressed in neural tissue. We found that the expression levels of *RBFOX1*, *RBFOX2*, *SRRM4*, *NOVA1*, and *PTBP1* all had high correlations (R^2^ > 0.35, FDR ≤ 0.05) to AS alterations in CTX (Figure 2e), but not in cerebellum (Figure 2f). Furthermore, enrichment analysis revealed that most changes in cortical AS occur in neuron-specific exons that are excluded in ASD (exons with ΔPSI > 50% in neurons overlap with exons excluded in ASD CTX, fold enrichment = 4.1, P = 1.8x10^−7^, Extended Data Fig. 5d).

To validate a regulatory relationship between splicing factors and these events, we evaluated experimental data from knockout, overexpression, and knockdown experiments for Rbfox1^20^, SRRM4^21^, and PTBP1^22^, respectively. We found that exons regulated by each of these splicing factors were significantly enriched in the set of exons excluded in ASD (Figure 2g), while in contrast, there was no enrichment for targets of ESRP^23^, a splicing factor involved in epithelial cell differentiation but not expressed in CTX. This shows that alterations in three splicing factors dysregulated in ASD regulate AS of the neuron-specific exons whose inclusion is dysregulated in ASD in CTX and not cerebellum, indicating selective alteration of neuronal splicing in ASD CTX. Remarkably, the expression patterns of these three splicing factors (and others for which appropriate validation experiments were unavailable) results in distinct clusters (Extended Data Fig. 5e), suggesting that subsets of splicing factors act in different individuals to mediate a common downstream AS alteration.

Taken together these results indicate global transcriptional alterations in ASD cerebral cortex, but not cerebellum at the level of protein coding transcripts, lncRNA and AS. Therefore, to determine how these different transcriptomic subcategories relate to each other in ASD, we compared the 1^st^ PC for each type of transcriptomic alteration across individuals (Figure 2h). Remarkably, the PCs are highly correlated (R^2^ > 0.8) indicating that the transcriptomic alteration is a unitary phenomenon across protein coding, noncoding, and splicing levels, rather than distinct forms of molecular alteration.

Previous analysis with gene expression microarrays in a small cohort suggested that the typical pattern of transcriptional differences between the frontal and temporal cortex may be attenuated in ASD^7^. To further test this possibility, we evaluated DGE between CTX regions (Supplementary Information) in 16 matched frontal and temporal CTX sample pairs from ASD and CTL subjects and found 551 genes differentially expressed between regions in controls, but only 51 in ASD (FDR ≤ 0.05; Figure 3a). We refer to the set of 523 genes with this pattern in CTL, but not ASD as the “Attenuated Cortical Patterning” set. The attenuation of patterning is evident from the global distribution of test statistics between frontal and temporal CTX in ASD and CTL and genes in this set do not show a greater difference in variability in ASD versus controls compared to other genes (Kolmogorov-Smirnov test, two-tailed P = 0.11, Extended Data Fig. 6a).

**Figure 3.**
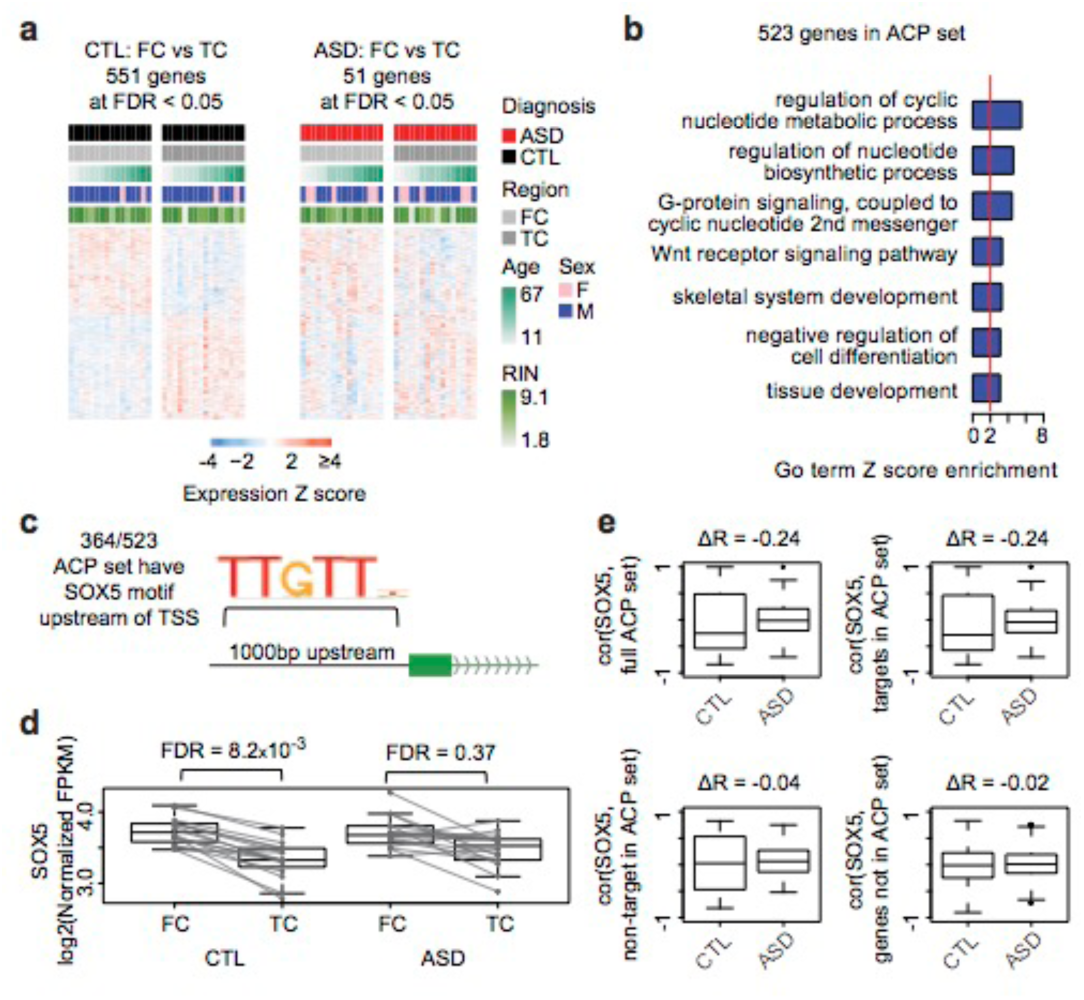
Attenuation of cortical patterning in ASD cortex. a, Heatmap of 551 genes exhibiting cortical patterning between frontal cortex (FC) and temporal cortex (TC) in ASD, with samples sorted by diagnostic status and brain region. b, Gene ontology term enrichment analysis of genes exhibiting attenuated cortical patterning (ACP). c, Schematic of transcription factor motif enrichment upstream of genes in the ACP set, with the *SOX5* motif sequence logo. d, The *SOX5* gene exhibits attenuated cortical patterning in ASD CTX compared to CTLs. Lines connect FC-TC pairs that are from the same individual. e, Correlation between *SOX5* gene expression and predicted targets in CTL and ASD, with all ACP genes (top left), SOX5 targets from the ACP set (top right), SOX5 non-targets from the ACP set (bottom left), and all genes not in the ACP set (bottom right). Plots show the difference in correlation between *SOX5* and other genes in ASD and CTL (ΔR).

We complemented this analysis with a machine learning approach using all 123 cortical samples, training a regularized regression model^24^ to classify frontal versus temporal CTX with independent gene expression data from BrainSpan^25^ (Extended Data Fig. 6b, Supplementary Information). Multiple approaches to training the classifier with BrainSpan can differentiate between frontal and temporal CTX in both CTL and ASD (Extended Data Fig. 6c-e), demonstrating that dissection and sample quality in our samples are of high quality. Loss of classification accuracy in ASD compared to CTL was observed when restricting the model to the genes with the most attenuated patterning in ASD (Extended Data Fig. 6f), demonstrating that attenuation of patterning generalizes across all samples. The Attenuated Cortical Patterning set includes multiple genes known to be involved in cell-cell communication and cortical patterning, such as *PCDH10*, *PCDH17*, *CDH12, MET*, and *PDGFD*, which was recently shown to mediate human specific aspects of cerebral cortical development^26^. GO term enrichment analysis of the Attenuated Cortical Patterning set identified enrichment for G protein coupled signaling, Wnt receptor signaling, and calcium binding, among several developmental processes (Figure 3b), and cell type enrichment analysis did not identify a strong preference for a particular cell type (Extended Data Fig. 6g).

To identify potential drivers of the alteration in cortical patterning, we evaluated transcription factor binding site enrichment upstream of genes in the Attenuated Cortical Patterning set (Supplementary Information), and found an enrichment of *SOX5* binding motifs (upstream of 364/523 genes, Figure 3c). Remarkably, *SOX5* itself belongs to the Attenuated Cortical Patterning set: while *SOX5* is differentially expressed between frontal and temporal CTX in CTL, it is not in ASD (Figure 3d). We thus predicted that if *SOX5* regulates cortically patterned genes, its expression should correlate with target gene transcript levels. Consistent with this prediction, we found that genes in the Attenuated Cortical Patterning set are anti-correlated with *SOX5* in CTL CTX, but not in ASD CTX (Figure 3e, top left; Wilcoxon rank sum test of R values, P = 0.01), suggesting that the normal role of SOX5 as a transcriptional repressor may be disrupted in ASD. We reasoned that a true loss of SOX5-mediated cortical patterning would be specific to the predicted SOX5 targets. Consistent with this, we find a loss of correlations between *SOX5* and predicted targets, but no difference in correlations between *SOX5* and non-targets in the Attenuated Cortical Patterning set (Figure 3e). Taken together, these findings show that a loss of regional patterning downstream of the transcriptional repressor *SOX5*, which plays a crucial role in glutamatergic neuron development^27,28^, contributes to the loss of regional identity in ASD.

Gene expression changes in postmortem brain may be a consequence of genetic factors, environmental factors, or both. Brain tissue from individuals with ASD that harbor known, penetrant genetic causes are very rare. However, we were able to identify postmortem brain tissue from 8 subjects with one of the more common recurrent forms of ASD, Duplication 15q Syndrome (dup15q, which is present in about 0.5-1% of ASD cases, see Extended Data Fig. 7a for characterization of duplications). We performed RNA-seq across frontal and temporal cortex and compared DGE changes in dup15q with those observed in individuals with idiopathic ASD to better understand the extent to which the observed molecular pathology overlaps. As expected, most genes in the 15q11.1-13.2 duplicated region have higher expression in dup15q CTX compared to CTL (Figure 4a), although *SNRPN* and *SNURF* were notably downregulated. Conversely, no significant upregulation of genes in this region were identified in idiopathic ASD or controls. Strikingly, when we assessed genome-wide expression changes, we observed a strong signal of DGE in dup15q that widely overlaps with that of idiopathic ASD (fold changes at FDR ≤ 0.05 in dup15q correlate with idiopathic ASD with R^2^ = 0.79, Figure 4b). Moreover, the slope of the best-fit line through these changes is 2.0, indicating that on average, the transcriptional changes in dup15q CTX are highly similar, but twice the magnitude of those observed in ASD CTX.

**Figure 4.**
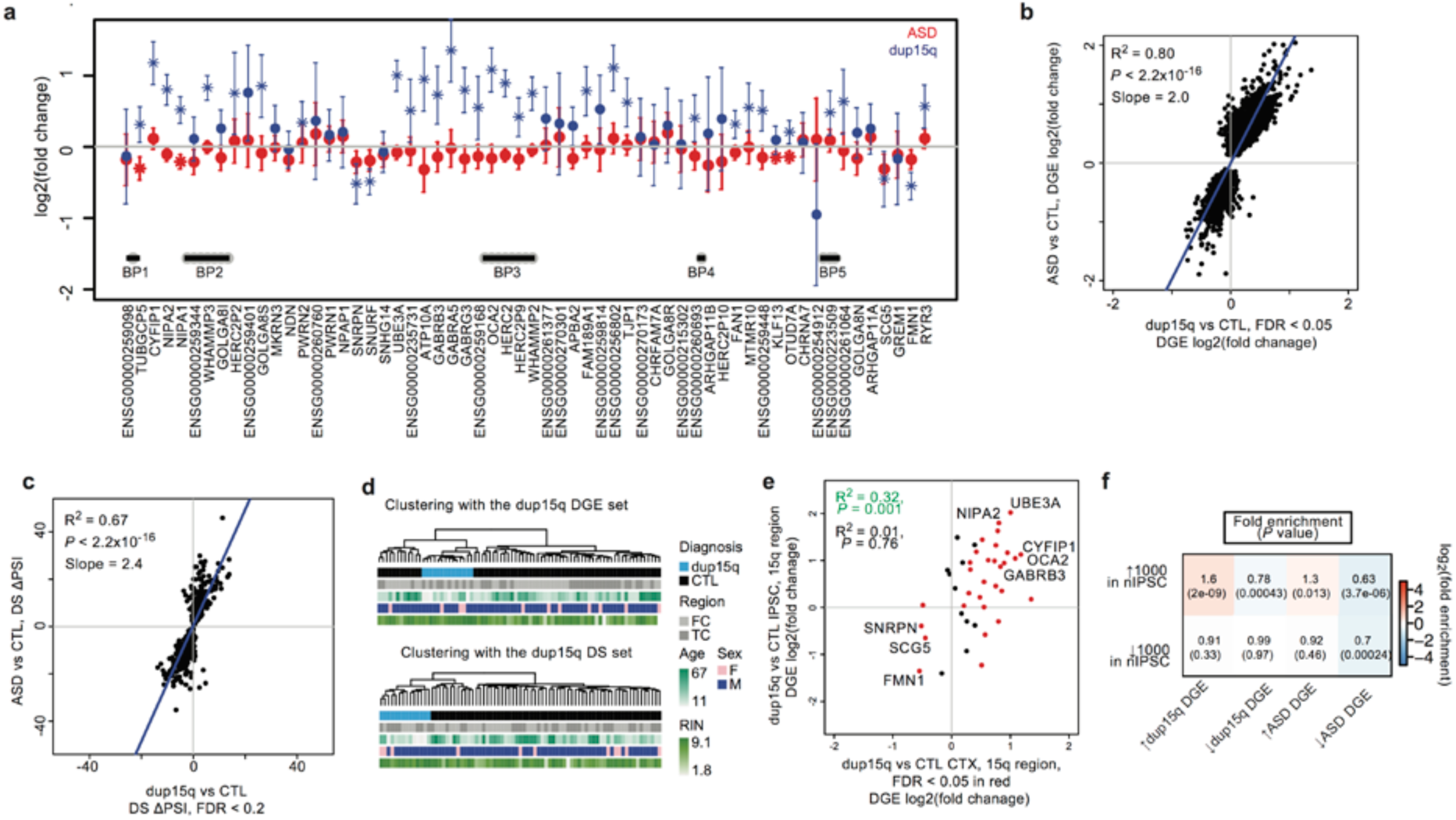
Duplication 15q Syndrome recapitulates transcriptomic changes in idiopathic ASD. a, DGE changes across the 15q11-13.2 region for ASD and dup15q compared to CTL, error bars are +/- 95% confidence intervals for the fold changes. b, Comparison of effect sizes in dup15q vs CTL and ASD vs CTL, with changes in dup15q at FDR ≤ 0.05 highlighted. c, Comparison of differential splicing (DS) changes in dup15q vs CTL and ASD vs CTL, highlighting 402 events at FDR ≤ 0.2 in dup15q. d, Average linkage hierarchical clustering of dup15q samples and controls using the DGE and DS gene sets. e, Plot of fold changes between induced pluripotent stem cells differentiated into neurons (nIPSCs) from dup15q vs CTL and postmortem CTX DGE from dup15q vs CTL in the 15q region. f, Heatmap overlapping the top 1000 genes up- and down- regulated in the nIPSC comparison to the up- and down- regulated genes in dup15q and idiopathic ASD CTX.

Next, we sought to evaluate AS changes in dup15q. There is only one significant splicing change in the dup15q region (Supplementary Table 3), consistent with the idea that duplication in this region duplicates all isoforms of the genes, resulting in minimal alteration of transcript structure. Similar to DGE, global AS analysis in dup15q CTX vs to CTL CTX revealed a stronger, but highly overlapping signature with idiopathic ASD CTX (fold changes at FDR ≤ 0.2 in dup15q agree correlate with idiopathic ASD with R^2^ = 0.66) indicating that splicing changes in dup15q syndrome recapitulate those of idiopathic ASD (Figure 4c). The slope of the best-fit line through the PSI for spliced exons in dup15q CTX compared to those in ASD CTX is 2.5 similar to DGE. Notably, both gene expression and AS changes in dup15q implicated similar pathways as those found in idiopathic ASD (Extended Data Fig. 7c-d). Clustering dup15q samples and CTL samples using both the DGE set and the differential AS set showed that all dup15q samples cluster together (Figure 4d), as opposed to the more variable clustering of idiopathic ASD, supporting the hypothesis that this shared genetic abnormality leads to a more homogeneous molecular phenotype.

Next, to test whether this molecular ASD signature may be due to independent of postmortem or reactive effects (Supplementary Information), we compared our data with gene expression profiles from a iPSC-derived neurons (nIPSCs)^29^ from dup15q were available, we could use these data to definitively reveal which changes in dup15q CTX are independent of postmortem or reactive effects (Supplementary Information), since such effects are not present *in vitro*. We observe that DGE in the 15q region is concordant with that seen in the nIPSCs (Figure 4e), even though the sample size is small and the analysis is likely underpowered. Upregulated changes in dup15q are also seen in nIPSCs (Figure 4f), consistent with our other statistical analyses showing limited effects of potential confounders. The very immature, fetal state of the nIPSCs^30^ likely explains the absence of an enrichment signal for genes downregulated in postnatal ASD brain, which are enriched for genes involved in neurons with more mature synapses.

We next applied gene network analysis to construct an organizing framework to understand shared biological functions across idiopathic ASD and dup15q (combining the ASD Discovery Set, ASD Replication Set, and dup15q set). We utilized Weighted Gene Co-expression Network Analysis (WGCNA), which identifies groups of genes with shared expression patterns across samples (modules) from which shared biological function is inferred. Modules identified via WGCNA can than be related to a range of relevant phenotypes and potential confounders^31,32^. We applied signed co-expression analysis and used bootstrapping to ensure the network was robust, and not dependent on any subset of samples (Supplementary Information), while controlling for technical factors and RNA quality (“Adjusted FPKM” levels, Methods). WGCNA identified 16 co-expression modules (Extended Data Fig. 8a, Supplementary Table 2), which are further characterized by their association to ASD (Extended Data Fig. 8b), enrichment for cell-type specific genes (Extended Data Fig. 8c), and enrichment for GO terms (Extended Data Fig. 9). Of the downregulated modules, three are associated with ASD and dup15q (M1/10/17) and one with dup15q only (M11). Five of the upregulated modules are associated with ASD and dup15q (M4/5/6/9/12) and one is specific to dup15q (M13) (Figure 5a, top). Additionally, we identified a module strongly enriched for genes from the Attenuated Cortical Patterning set and *Wnt* signaling that contains *SOX5* (M12; fold enrichment = 3.0, P = 3x10^−8^), verifying the strong relationship observed between the *Wnt* pathway regulating TF *SOX5* and attenuation of cortical patterning^33^.

**Figure 5.**
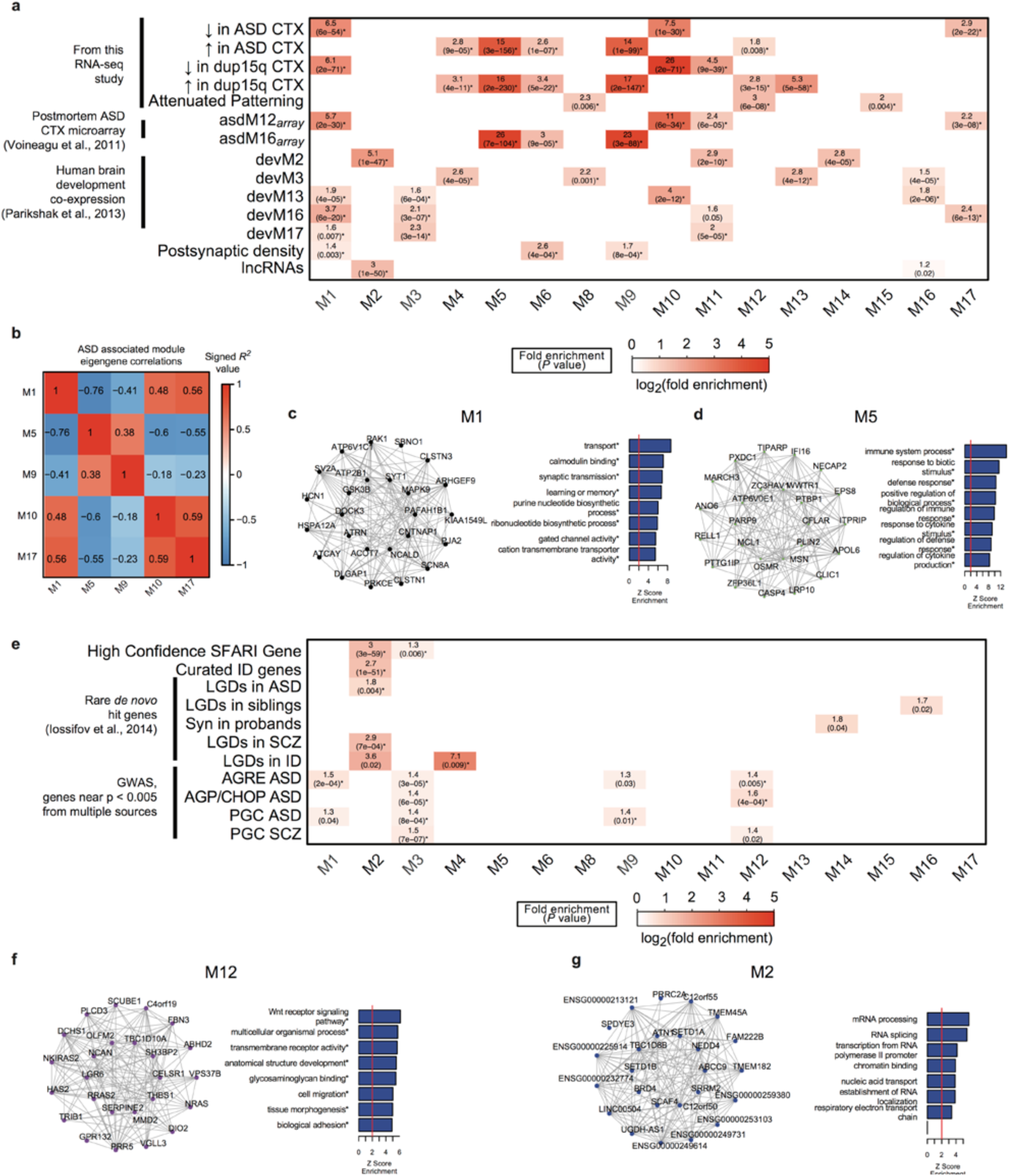
Co-expression network analysis across all ASD and CTL samples in CTX. a, Gene set enrichment analyses comparing the 16 co-expression modules with multiple gene sets from this RNA-seq study, from postmortem ASD CTX microarray, from human brain development, from the postsynaptic density and set of all brain-expressed lncRNAs. b, Comparison of five ASD-associated modules against each other by correlating module eigengenes. c, Module plot of M1 displaying the top 25 hub genes along with the module’s Gene Ontology term enrichment. d, similar to c, but for M5. e, Gene set enrichment analysis with genome-wide whole-exome sequencing data (Rare *de novo* hit genes) and genome-wide association study (GWAS) results in ASD, schizophrenia (SCZ), and intellectual disability (ID). Boxes are filled if the odds ratio is greater than 0, and the enrichment *P* < 0.05. Asterisks* indicate FDR ≤ 0.05 across all comparisons in a and e. f,g, similar to c, but for M12 and M2, respectively. Abbreviations: LGD, likely gene disrupting, genes affected by nonsense, nonsynonymous, or splice-site mutations or frame-shift indels; AGRE, AGP/CHOP, and PGC refer to consortia that collect genetic data (Supplementary Information for details).

Notably, the modules identified here significantly overlap with previous patterns identified in ASD (asdM12_*array*_ and asdM16_*array*_^7^; Figure 5a, middle). We found that the ASD-associated modules identified by our larger sample size and RNA-seq provide significant refinement of previous observations by identifying more discrete biological processes related to cortical development^34^, the post-synaptic density^35^, and lncRNAs (Figure 5a, bottom). For example, M1 overlaps a subset of asdM12_*array*_ (fold enrichment = 5.7) and developmental modules (devM16 fold enrichment = 3.7), and is enriched for proteins found in the PSD and genes involved in calcium signaling and gated ion channel signaling. Another subset of asdM12_*array*_, M10 (fold enrichment = 11) overlaps more with a mid-fetal upregulated cortical development module (devM13 fold enrichment = 4.0), and genes involved in secretory pathways and intracellular signaling. A third module, M17 shows the least overlap with asdM12_*array*_ (fold enrichment = 2.2) and is related to energy metabolism. Notably, these three modules are enriched for neuron-specific genes (Extended Data Fig. 8c), but not all neuronal modules are down regulated in ASD (M3 is not altered in ASD CTX). Taken together, specific neurobiological processes are affected in individuals with ASD related to developmentally regulated neurodevelopmental processes.

The most upregulated modules, M5 and M9, both strongly overlap (fold enrichments > 20) with previously identified upregulated co-expression module asdM16_*array*_. M5 is enriched for microglial cell markers and immune response pathways, whereas M9 is enriched for astrocyte markers and immune-mediated signaling and immune cell activation (Extended Data Fig. 8c, Extended Data Fig. 9). This analysis clearly separates the contributions of the coordinated biological processes of microglial activation and reactive astrocytosis, which were previously not distinguishable as separate modules^7^. Thus, our analysis pinpoints more specific biological pathways in idiopathic ASD than those previously identified and reveals that similar changes occur downstream of the genetic perturbation in dup15q.

We evaluated the relationship between the five modules most strongly associated with ASD (M1/5/9/10/17, which are supported by module-trait association analysis and gene set enrichment analysis, Supplementary Information), and found that there was a remarkably high anti-correlation between the eigengene of M5 and downregulated modules, particularly M1 (R^2^ = 0.76) (Figure 5b). M1 (Figure 5c) is downregulated in ASD and enriched for genes at the PSD and genes involved in synaptic transmission, while M5 (Figure 5d) is enriched for microglial genes and cytokine activation. This strong anti-correlation between microglial signaling and synaptic signaling in ASD and dup15q provides evidence in humans for dysregulation of microglia-mediated synaptic pruning, as previously suggested^36^.

Next, to determine the role of causal genetic variation, we evaluated enrichment of both rare genetic variants, focusing on genes affected by ASD associated gene disrupting (LGD) *de novo* mutations^37^, and common variants^38,39^. Genes within three modules, M1, M3, and M12, show enrichment for common variation signal for ASD (Figure 5e, Methods). Remarkably, M12 (Figure 5f), which is related to cortical patterning and Wnt signaling, also exhibit GWAS signal enrichment, providing the first evidence that risk conferred by common variation in ASD may affect regionalization of the cortex. Interestingly, M3 is significantly enriched for both schizophrenia (SCZ) and ASD common variants, is related to synaptic transmission, nervous system development, and regulation of ion channel activity (Extended Data Fig. 9), consistent with the notion that ASD and SCZ share common and rare genetic risk^1,40-43^.

We only identified one module, M2 (Figure 5g), as significantly enriched in protein disrupting (nonsense, splice site, or frameshift) rare *de novo* variants previously associated with SCZ and ASD. M2 overlaps with a cortical developmental module implicated in ASD^34^ (devM2 fold enrichment = 5.1). Notably, M2 is not differential between ASD and CTL in our dataset, consistent with the observation that these genes are primarily expressed during early neuronal development in fetal brain^34^. Remarkably, M2 contains an unusually large fraction of lncRNAs (15% of the genes in M2 are classified as lncRNAs, while other modules are 1-5% lncRNA). We hypothesize that, in addition to protein coding genes involved in transcriptional and chromatin regulation, rare *de novo* variants may also affect lncRNAs in ASD, a prediction that will be testable once large sets of whole genome sequences are available.

These combined transcriptomic and genetic analyses reveal that different forms of genetic variation affect biological processes involved in multiple stages of cortical development. Common genetic risk is enriched in M3, M1, and M12, which reflect early glutamatergic neurogenesis, later neuronal function, and cortical patterning, respectively. We also observe that rare *de novo* variation, which is enriched in M2, affects distinct biology related to transcriptional regulation and chromatin modification. These findings are consistent with transcriptomic analyses of early prenatal brain development and ASD risk mutations that implicate chromatin regulation and glutamatergic neuron development^34,44^.

We provide the first comprehensive picture of largely unexplored aspects of transcription in ASD, lncRNA and alternative splicing, and identify a strong convergent signal in these, as well as protein coding genes^7^. These results will aid in interpreting genetic variation outside of the known exome, as whole genome sequencing supplants current methods. A role of lncRNAs has been previously explored in ASD^45^, but only two individuals were evaluated with targeted microarrays. We evaluate lncRNAs in an unbiased manner across many individuals, notably identifying an enrichment of lncRNAs in M2, most of which are uncharacterized in brain and arose on the primate lineage. The involvement of lncRNAs in this early developmental program that is enriched for *de novo* mutations implicated in ASD suggests their study will be particularly relevant to understanding the emergence of primate higher cognition on the mammalian lineage, and by extension human brain evolution^15,46,47^.

We also provide the first confirmation of an attenuation of genes that typically show differential expression between frontal and temporal lobe in ASD CTX and further identified *SOX5*, known to regulate cortical laminar development^50,51^, as a putative regulator of this disruption. That M12, which is enriched for genes exhibiting cortical regionalization and is also enriched in ASD GWAS signal, supports the prediction that attenuation of patterning may be mediated by common genetic variation in or near the *SOX5* target genes. Disruption of cortical lamination by direct effects on glutamatergic neurogenesis and function has been predicted by independent data, including network analyses of rare ASD associated variants identified in exome sequencing studies^34,44^.

These data, in conjunction with previous studies, reveal a consistent picture of the ASD’s emerging postnatal and adult pathology. Specific neuronal signaling and synaptic molecules are downregulated and astrocyte and microglial genes are upregulated in over 2/3 of cases. Microglial infiltration has been observed in ASD cortex with independent methods^52^, and normal microglial pruning has been shown to be necessary for brain development^36^. Our findings further suggest that aberrant microglial-neuronal interactions may be pervasive in ASD and related to the gene expression signature seen in a majority of individuals. In our comprehensive AS analysis, we identify three splicing factors upstream of the altered splicing signature observed in ASD CTX. These factors are known to be involved in coordinating sequential processes in neuronal development^17,21^ and maintaining neuronal function^48,49^. It may therefore be sufficient to disrupt any one of these factors to induce a similar outcome during brain development, which would be consistent with the shared downstream perturbation observed here.

Finally, evaluation of the transcriptome in dup15q supports the enormous value of the “genotype first” approach of studying syndromic forms of ASD, with known penetrant genetic lesions^53^. It is highly unlikely that the shared transcriptional dysregulation in dup15q is due to a shared environmental insult. Thus, the most parsimonious explanation for the convergent transcriptomic pathology seen in all dup15q and over 2/3 of the cases of idiopathic ASD is that it represents an adaptive or maladaptive response to a primary genetic insult, which in most cases of ASD will be genetic^2,54^. As future investigations pursue the full range of causal genetic variation contributing to ASD risk, these analyses and data will be valuable for interpreting genetic and epigenetic studies of ASD as well as those of other neuropsychiatric disorders.

## Methods

Sample description: Brain tissue for ASD and control individuals was acquired from the Autism Tissue Program (ATP) brain bank at the Harvard Brain and Tissue Bank and the University of Maryland Brain and Tissue Bank (a Brain and Tissue Repository of the NIH NeuroBioBank). Sample acquisition protocols were followed for each brain bank, and samples were de-identified prior to acquisition. Brain sample and individual level metadata is available in Supplementary Table 1.

RNA-seq methodology: Starting with 1ug of total RNA, samples were rRNA depleted (RiboZero Gold, Illumina) and libraries were prepared using the TruSeq v2 kit (Illumina) to construct unstranded libraries with a mean fragment size of 150bp (range 100-300bp) that underwent 50bp paired end sequencing on an Illumina HiSeq 2000 or 2500 machine. Paired-end reads were mapped to hg19 using Gencode v18 annotations^55^ via Tophat2^56^. Gene expression levels were quantified using union exon models with HTSeq^57^. For additional and information on sequencing and read alignment parameters, please see Supplementary Information.

Sample sets for analysis: For differential gene expression and splicing analysis, we defined an age matched set, referred to as the ASD Discovery Set (106 samples in CTX, 51 in cerebellum) of idiopathic ASD and control samples for the discovery set, and held out younger or unmatched samples as the ASD Discovery Set (17 in CTX, 8 in cerebellum). Dup15q individuals were analysed separately, utilizing the full set of controls from the ASD Discovery Set. For co-expression network analysis, we combined the discovery set, replication set, and dup15q individuals for a total of 137 CTX samples and 59 cerebellum samples.

Differential Gene Expression (DGE): DGE analysis was performed with expression levels adjusted for gene length, library size, and G+C content (referred to as “Normalized FPKM”) Supplementary Information. CTX samples (frontal and temporal) were analyzed separately from cerebellum samples. A linear mixed effects model framework was used to assess differential expression in log2(Normalized FPKM) values for each gene for cortical regions (as multiple brain regions were available from the same individuals) and a linear model was used for cerebellum (where one brain region was available in each individual, with a handful of technical replicates removed). Individual brain ID was treated as a random effect, while age, sex, brain region (except in the case of cerebellum, where there is only one region), and diagnoses were treated as fixed effects. We also used technical covariates accounting for RNA quality, library preparation, and batch effects as fixed effects into this model (Supplementary Information).

Reproducibility analyses: We assessed replication between datasets by evaluating the concordance between independent sample sets by comparing the squared correlation (R^2^) of fold changes of genes in each sample set at a non-stringent P value threshold. This general approach has been shown to be effective for identifying reproducible gene expression patterns^58^, and we modify it such that the P value threshold is set in one sample set (the *x* axis in the scatterplots), and the R^2^ with fold changes in these genes are evaluated in an independent sample set (the *y* axis in the scatterplots).

Differential Splicing Analysis: Alternative splicing was quantified using the percent spliced in (PSI) metric using Multivariate Analysis of Transcript Splicing (MATS, v3.08)^18^. For each event, MATS reports counts supporting the inclusion (I) or exclusion (E) of a splicing event. To reduce spurious events due to low counts, we required at least 80% of samples to have I + S >= 10. For these events, the percent spliced in is calculated as PSI = I / (I + S) (Extended Data Fig. 4a). Statistical analysis for differential splicing was performed utilizing the linear mixed effects model regression framework as described above for DGE. This approach is advantageous over existing methods as it allows modeling of covariates and takes into consideration the variability in PSI across samples when assessing event significance with ASD (Supplementary Information).

Genotyping dup15q: For Dup15q samples, the type of duplication and copy number in the breakpoint 2-3 region were available for these brains^59^. To expand this to the regions between each of the recurrent breakpoint in these samples, 7/8 dup15q brains were genotyped (one was not genotyped due to limitations in tissue availability). The number of copies between each of the breakpoints is reported in Extended Data Fig. 7a.

Co-expression network analysis: The R package weighted gene co-expression network analysis (WGCNA) was used to construct co-expression networks using the technical variation normalized data^31,60^ (referred to as “Adjusted FPKM”). We used the biweight midcorrelation to assess correlations between log2(Normalized FPKM) and parameters for network analysis are described in Supplementary Information. Notably, we utilized a modified version of WGCNA that involves bootstrapping the underlying dataset 100 times and constructing 100 networks. The consensus of these networks (50^th^ percentile across all edges) was then used as the final network ^32^, ensuring that a handful of samples do not determine the network structure. For module-trait analyses, 1^st^ principal component of each module (eigengene) was related to ASD diagnosis, age, sex, and brain region in a linear mixed effects framework as above, only replacing the expression values of each gene with the eigengene.

Enrichment analysis of gene sets and GWAS: Enrichment analyses were performed either with Fisher’s exact test (cell type and splicing factor enrichments) or logistic regression (all enrichment analyses in Figure 5). We used logistic regression in the latter case to control for gene length or other biases that may influence enrichment analysis (Supplementary Information). All GO term enrichment analysis was performed using GO Elite^61^ with 10,000 permutations. We focused on molecular function and biological process terms for display purposes.

## Extended Data Figures & Legends

**Extended Data Figure 1.**
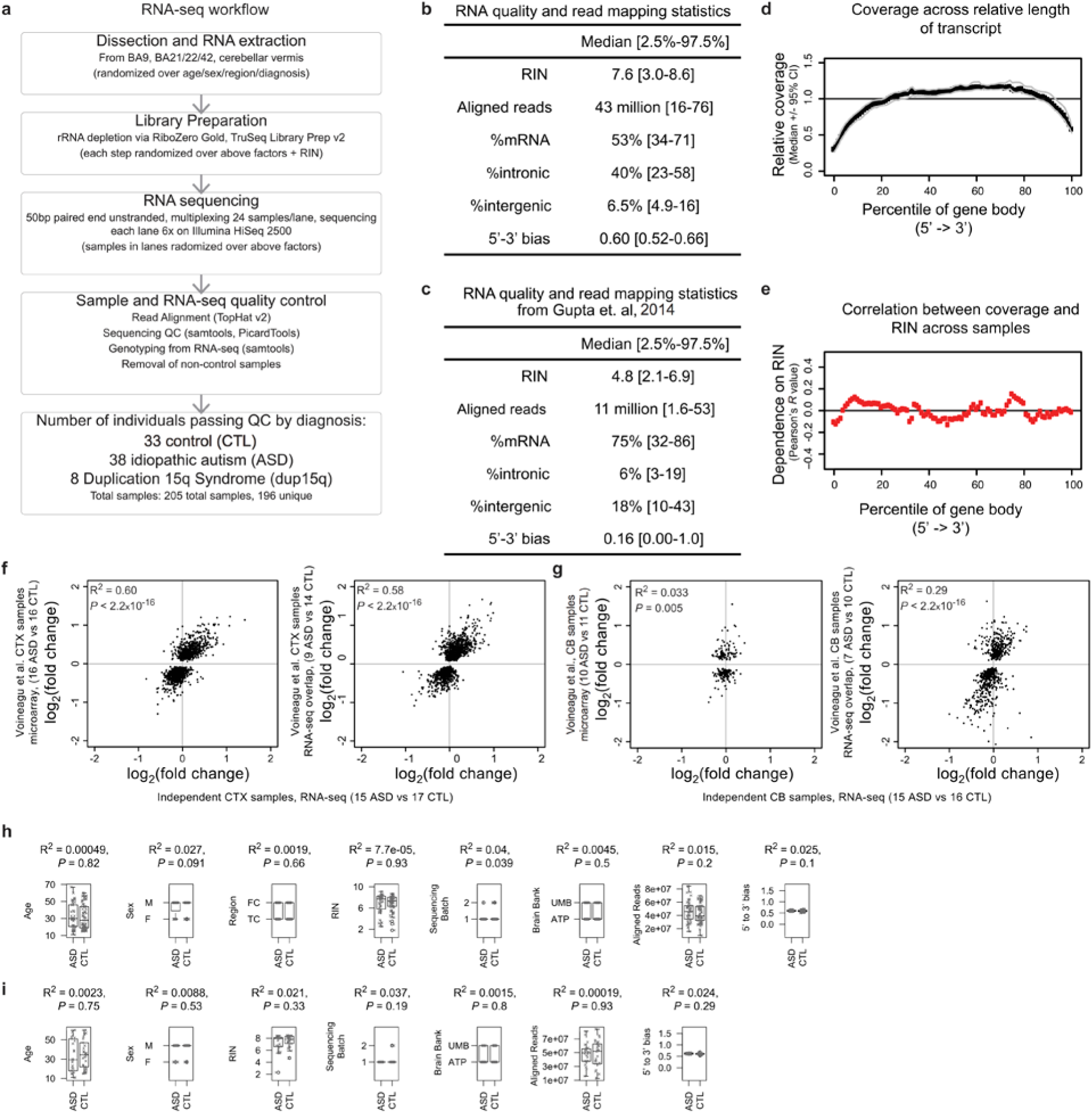
Methodology, quality control, and differential expression replication analysis. a, RNA-seq workflow, including RNA extraction, library preparation, sequencing, read alignment, and quality control. b, RNA-seq quality and alignment statistics from this study, including RNA integrity number (RIN), number of aligned reads, proportion of reads mapping to different genomic features (mRNA, intronic, intergenic), and bias in coverage from the 5’ to the 3’ end of the top 1000 expressed transcripts (statistics compiled using PicardTools). c, Similar statistics as in b for another RNA-seq study that utilized polyA tail selection mRNA-seq to evaluate the transcriptome in ASD cortex^11^ (primarily BA19, visual cortex, but also including some BA10/44 samples, frontal cortex). d, RNA-seq read coverage relative to normalized gene length across transcripts from the 5’ to the 3’ end in this study. e, Dependence between coverage and RIN across gene body (correlation between RIN and coverage in d across samples). f, Correlation of ASD vs CTL fold changes between previously evaluated and new ASD samples in CTX by microarray (left) and RNA-seq (right) using genes that were at *P* < 0.05 the samples from Voineagu et al., 2011. g, Correlation between effect sizes as in f, but for cerebellum (CB) samples. h,i, Correlation between covariates and ASD vs CTL status in CTX (h) and CB (i) in the ASD Discovery Set.

**Extended Data Figure 2.**
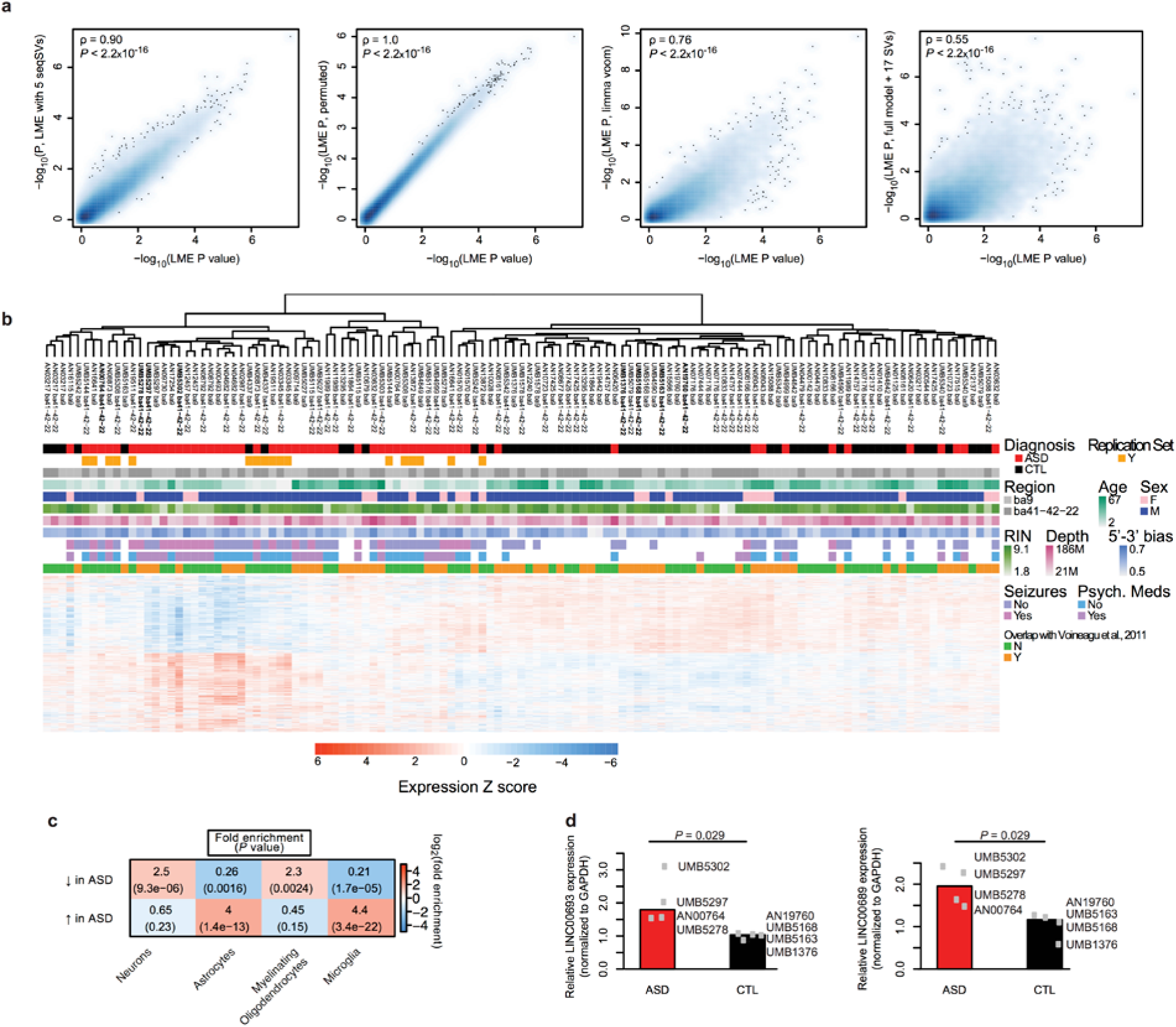
Transcriptome-wide differential gene expression (DGE) analysis in CTX. a, Comparison of P value rankings across different methods for DGE with Spearman’s correlation. From left to right: removal of three additional principal components of sequencing statistics (Supplementary Information) related to RNA-sequencing quality, application of a permutation analysis for DGE P value computation, application of variance-weighted linear regression for DGE^62^, and using surrogate variable analysis for DGE^63^. b, Average linkage hierarchical clustering heatmap using all genes DGE in the ASD Discovery Set, but including all idiopathic ASD frontal cortex (FC) and temporal cortex (TC) samples across 123 samples, combining the ASD Discovery set and the ASD Replication set. Bolded samples in the dendrogram are used for validation in d. c, Enrichment analysis of cell-type specific gene sets (5-fold enriched in the cell type compared to all other cells) with genes decreased and increased in ASD. d, RT-PCR validation of the two lincRNAs shown in Figure 1f-g, P values are computed with the Wilcoxon rank-sum test.

**Extended Data Figure 3.**
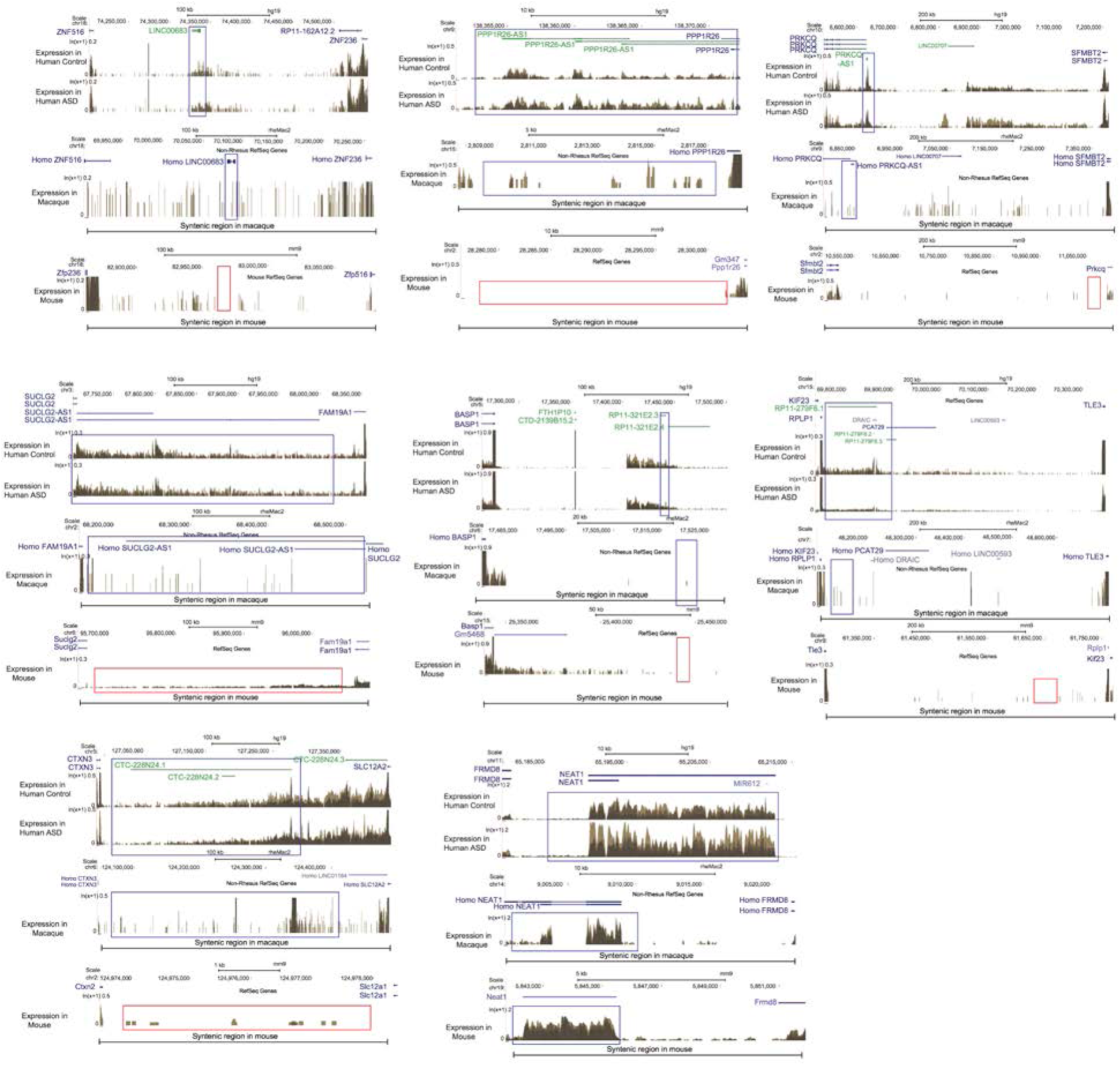
Gene browser tracks for selected primate-specific lncRNAs. For each lncRNA, expression for representative samples for ASD vs CTL (top) in human, macaque (middle), and mouse (bottom) are shown. The genome location for macaque and mouse displayed is syntenic to the human region, with the expected location of the lncRNA highlighted.

**Extended Data Figure 4.**
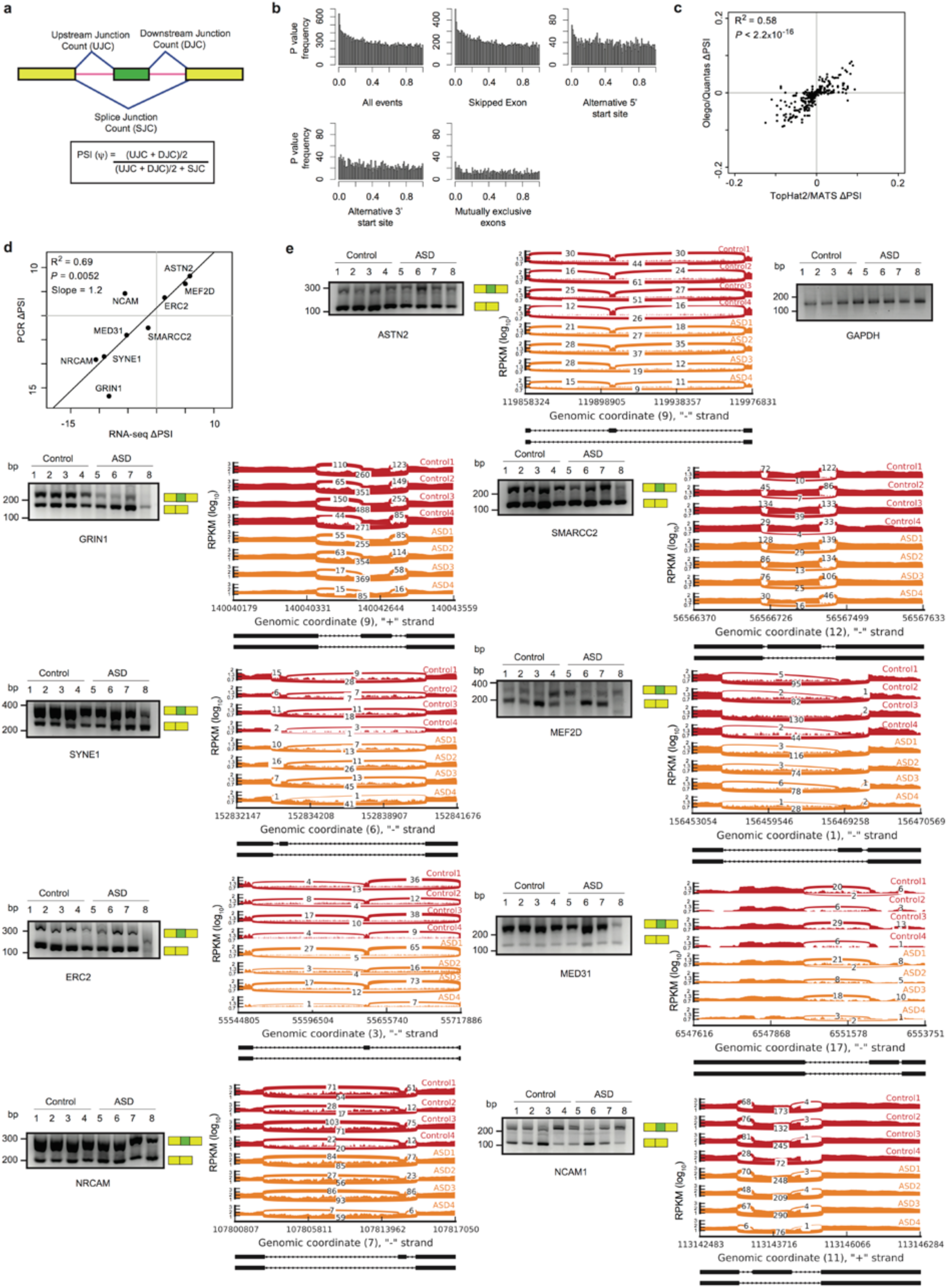
Splicing analyses and validation in ASD. a, Schematic describing how the percent spliced in (PSI) metric is computed. b, Distribution of *P* values for changes in the PSI between ASD and CTL in CTX for all events (left) and event subtypes (SE, spiced exon; A5SS, alternative 5’ splice site; A3SS, alternative 3’ splice site; MXE, mutually exclusive exons). c, Comparison of the CTX splicing analyses in when using PSI values obtained via read alignment by TopHat2^64^ followed by the MATS^18^ pipeline (used throughout this study) against read alignment by OLego followed by Quantas^65^. d, Comparison of ΔPSI values in nine splicing events between PCR and RNA-seq. e, PCR validation and sashimi plots for the nine splicing events delineated in d, from the samples highlighted in Extended Data Fig. 5a.

**Extended Data Figure 5.**
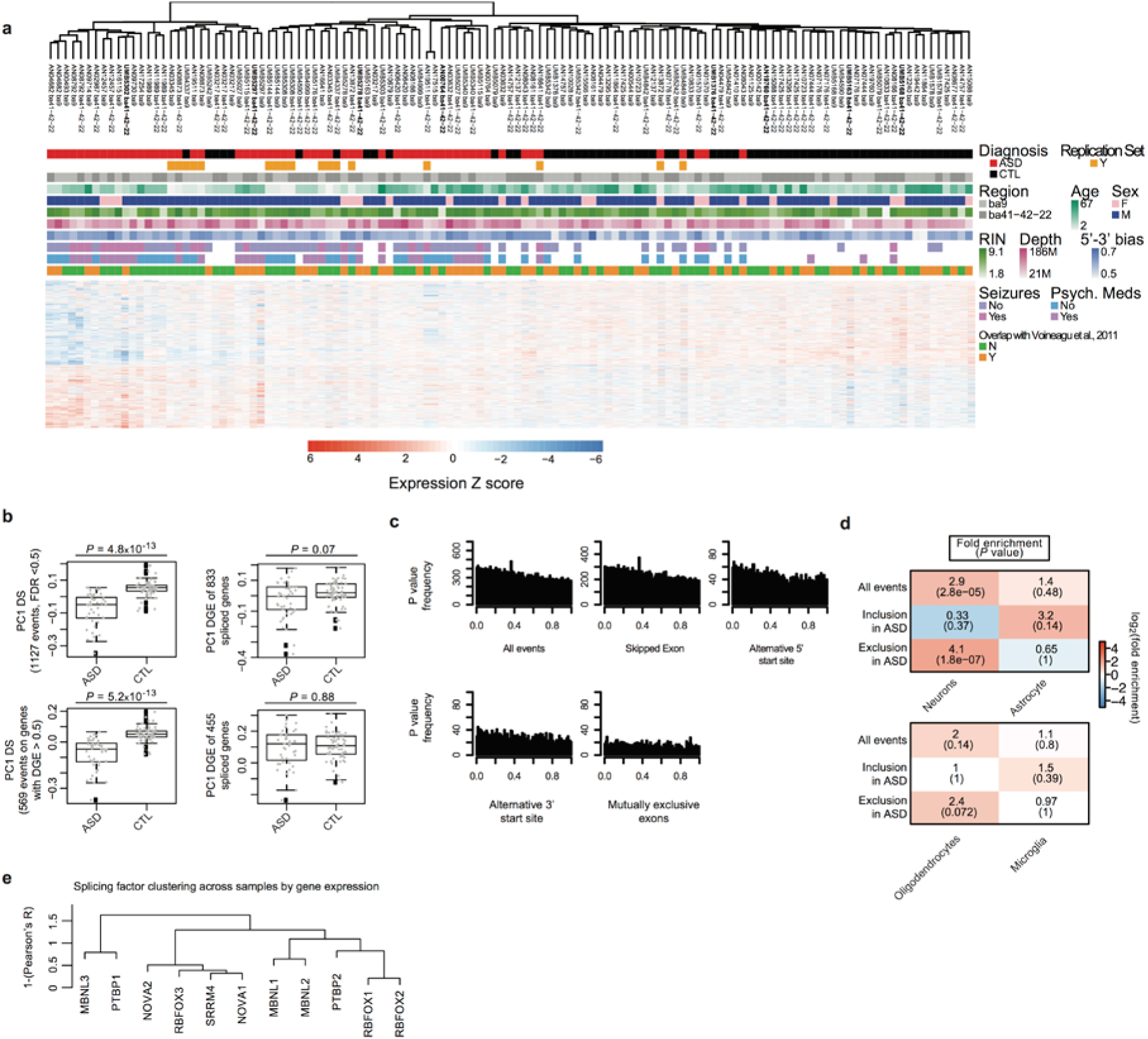
Additional splicing analyses in ASD. a, Average linkage hierarchical clustering heatmap using all differentially spiced (DS) events from the ASD Discovery Set, but including all idiopathic ASD neocortical samples (FC and TC) across 123 samples, combining the ASD Discovery set and the ASD Replication set. Bolded samples in the dendrogram were used for PCR validation in Extended Data Fig. 4. b, Top: difference between ASD and CTL in the DS set based on PC1 of the DS set at the PSI level, and PC1 of the gene expression levels of genes in the DS set. Bottom: Same comparison after differentially expressed genes (p < 0.05) are removed. c, Distribution of P values for changes in the PSI between ASD and CTL in cerebellum. d, Cell-type enrichment analysis of splicing events from CTX. e, Average-linkage hierarchical clustering using 1-(Pearson’s correlation) to compare the gene expression patterns of the splicing factors investigated in Figure 2.

**Extended Data Figure 6.**
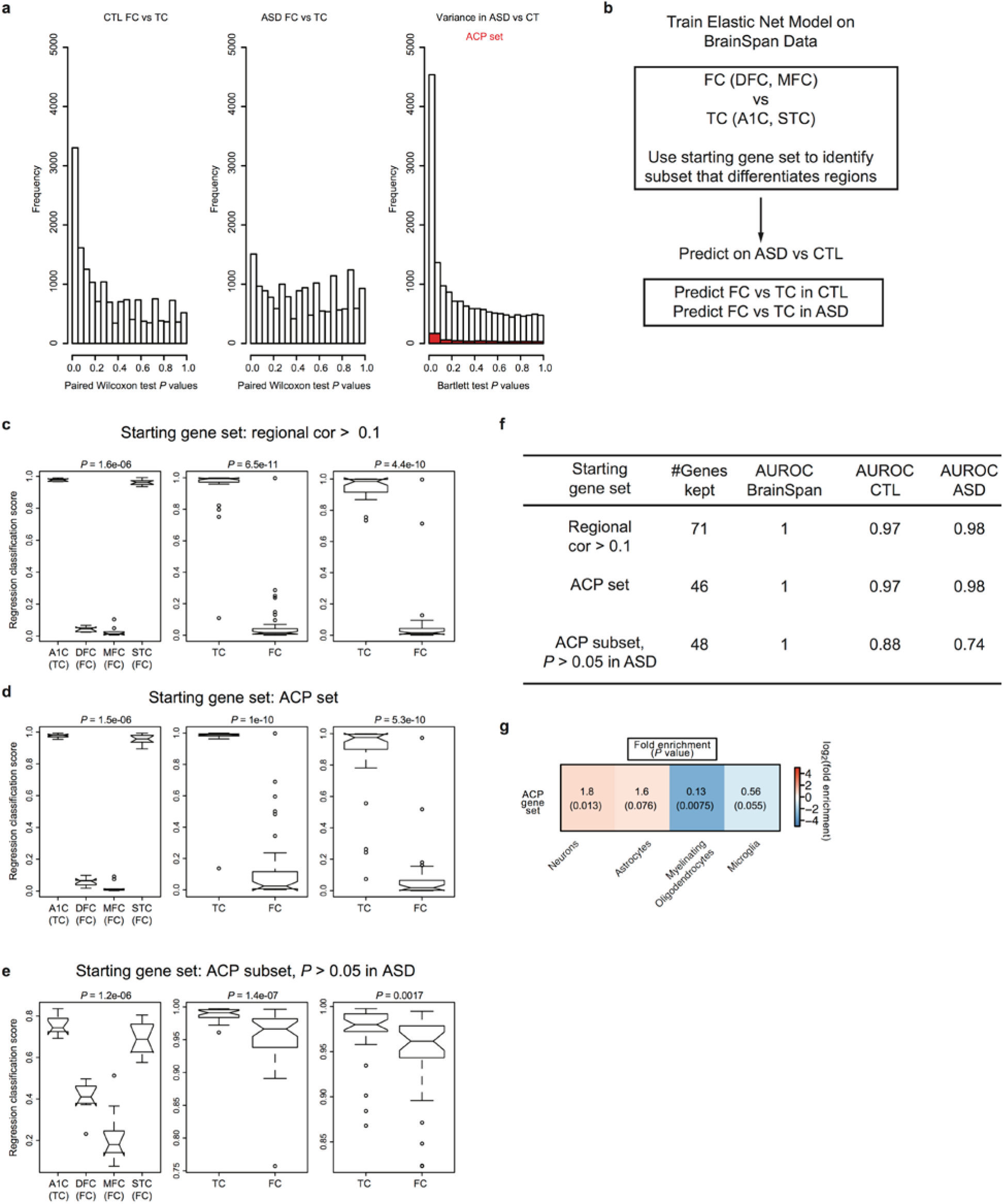
Attenuation of cortical patterning in ASD. a, Histograms of P values from paired Wilcoxon rank-sum test differential gene expression between 16 frontal cortex (FC) and 16 temporal cortex (TC) in CTL and ASD and a histogram of Bartlett’s test P values for differences in gene expression variance between ASD and CTL for all genes (white) and genes in the Attenuated Cortical Patterning (ACP) set (red). c, Approach to training the elastic net model on BrainSpan and application of the model on 123 cortical samples in this study. c-e, Results of learned cortical region classifications with different starting gene sets, with the BrainSpan training set (left), CTL samples (middle), and ASD samples (right) in each panel and the Wilcoxon rank-sum test P value of FC vs TC difference for each comparison. f, Summary of results form c-e. g, Cell type enrichment analysis for genes in the ACP set. Abbreviations: A1C, primary auditory cortex; DFC, dorsolateral prefrontal cortex; MFC, medial prefrontal cortex; STC, superior temporal cortex; FC, frontal cortex; TC, temporal cortex; AUROC, area under the receiver-operator characteristic curve.

**Extended Data Figure 7.**
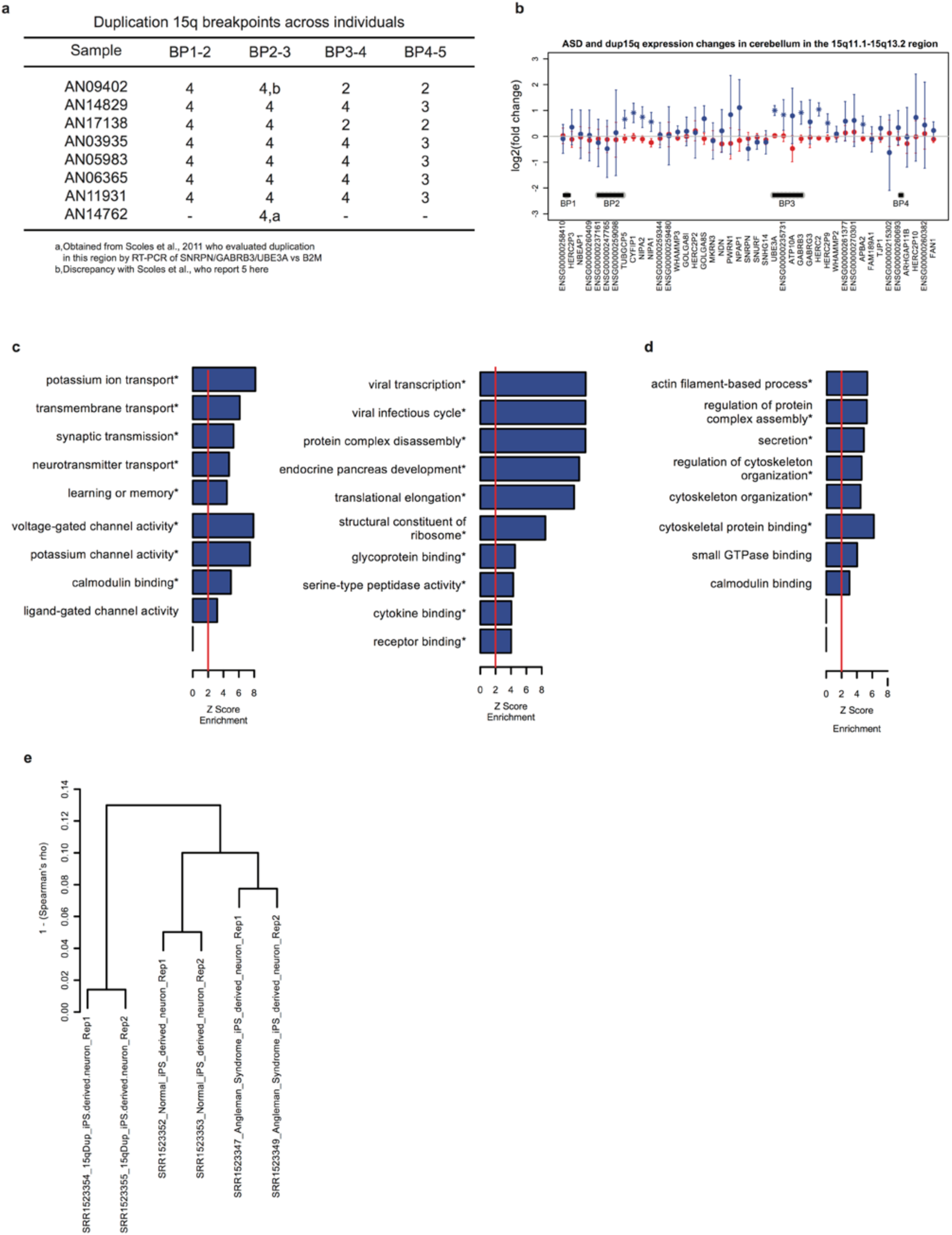
Dup15q syndrome analyses. a, Copy number between breakpoints (BP) in the 15q region. Genome-wide CNV analysis allowed evaluation of copy number in additional regions from previous studies^59,66^. b, Differential expression across the 15q region of interest in dup15q vs CTL and ASD vs CTL cerebellum, note only 3 samples were available for dup15q cerebellum so additional analyses were not pursued. c, Gene Ontology term enrichment analysis for the dup15q CTX differential expression set. d, Gene Ontology term enrichment analysis for the dup15q CTX differential splicing (DS) set. e, Hierarchical clustering of iPSC-derived neurons from dup15q, Angelman syndrome, and a control^29^.

**Extended Data Figure 8.**
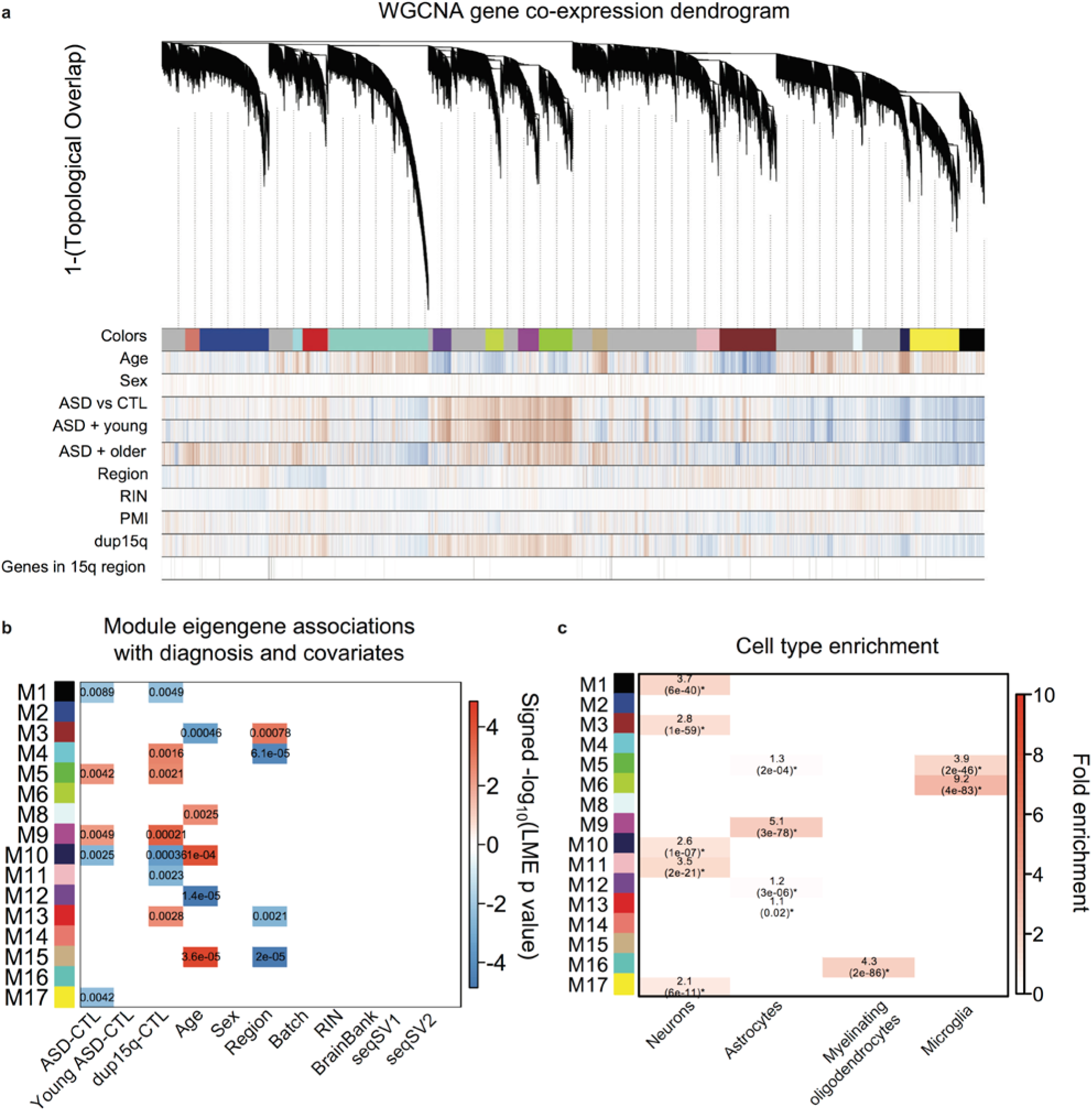
Co-expression network analysis in ASD CTX. a, Modules identified from a dendrogram constructed from a consensus of 100 bootstrapped datasets using the 137 CTX samples. Correlations for each gene to each measured factor are delineated below the dendrogram (blue = negative, red = positive correlation). b, Module-trait associations as computed by a linear mixed effects model with all factors on the x-axis used as covariates. All P values are displayed where the coefficient passed p < 0.01. Note that this alternative approach to module-trait association agrees with the Fisher’s exact test used in Figure 5a when the fold enrichment for module overlap with DGE sets is > 2.8, and we use an intersection of both methods for the modules focused on in Figure 5b. c, Module enrichments for cell type specific gene expression patterns.

**Extended Data Figure 9.**
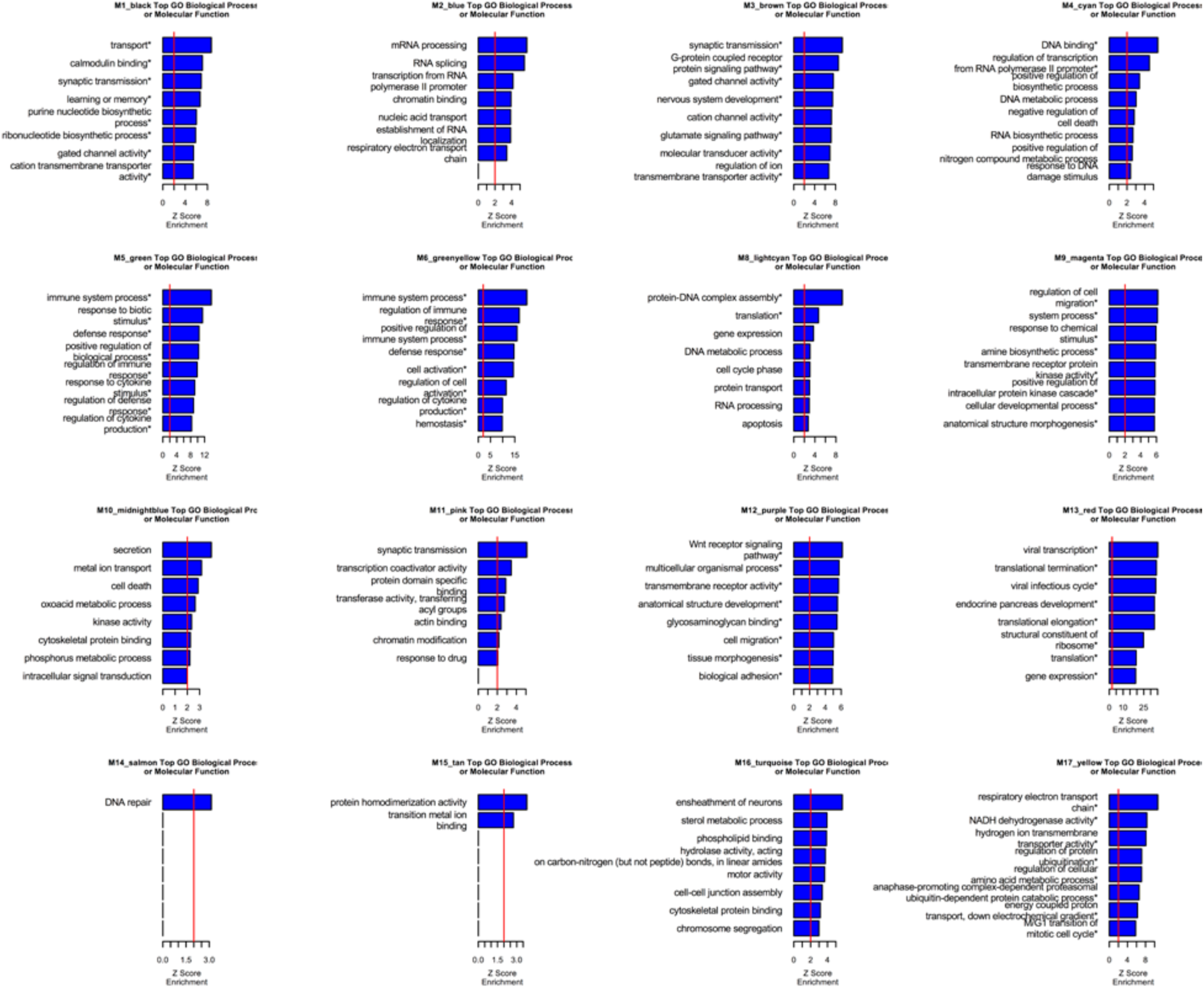
GO term enrichments for all modules. *FDR < 0.05 across all GO enrichments across all modules.

## End Notes

None.

## Author Contributions

NNP, VS, and TGB performed dissections and RNA-seq analyses and differential gene expression analysis. NNP and VS performed splicing and co-expression network analysis. NNP and TGB performed analyses with Duplication 15q Syndrome individuals. NNP and MG reviewed clinical information and performed meta-analysis of ASD gene expression studies. VL and JKL performed genotyping and CNV analysis on dup15q samples. VS performed validation experiments for gene and splicing level alterations in ASD. MI and BJB assisted with splicing analyses. RJ performed dissections. SH provided guidance on differential gene expression and co-expression analyses. DHG provided guidance on all experiments and analyses. NNP and DHG wrote the manuscript. All authors contributed to revising and finalizing the manuscript.

